# Sialic acids are a barrier to the entry of non-influenza orthomyxoviruses

**DOI:** 10.64898/2026.01.15.699645

**Authors:** Gabriel Dupré, Jorge Moreno-García, Elias Bendl, Georg Kochs, Jérémy Dufloo, Rafael Sanjuán

## Abstract

Sialic acids (SAs) are abundantly expressed on vertebrate cell surfaces and are widely recognized as key viral attachment factors, particularly for influenza viruses. However, their role remains understudied in other orthomyxoviruses, such as thogoto and quaranja viruses, which are tick-borne viruses sporadically infecting humans. Enzymatic removal of SAs increased the infectivity of Thogoto and Dhori viruses, as well as pseudotypes carrying the glycoproteins of Oz, Sinu, and Wellfleet Bay viruses. A similar effect on pseudotype infectivity was observed following the binding of specific lectins to SAs. These findings indicate that, in contrast to influenza viruses, SAs act as a barrier to the entry of these orthomyxoviruses. Experimental evolution of the Sinu and Wellfleet Bay virus glycoproteins revealed point mutations that partially overcame this barrier. Given the abundance of sialic acids in mucosal tissues, we speculate that SAs may contribute to the inability of thogoto and quaranjaviruses to transmit directly between vertebrate hosts. Our results also underscore the importance of monitoring the circulation of these viruses for potential changes in their transmission routes.

**Author summary:** Many viruses, including influenza viruses, use sugar molecules called sialic acids (SAs) on the surface of host cells to promote viral entry. Here, we show that SAs can instead restrict infection for several lesser-known relatives of influenza virus that are primarily transmitted by ticks and can occasionally infect humans. Removal of SAs from human cells strongly increased entry of these non-influenza viruses, when using both non-replicative pseudotypes and authentic viruses. In addition, masking SAs also enhanced entry of viral pseudotypes. We further found that viral surface proteins can acquire mutations that partially reduce this restriction, indicating an ability by these viruses to adapt to SA–rich environments. Because SAs are abundant in vertebrate mucosal tissues such as the respiratory tract, our findings suggest a possible explanation for why these viruses do not transmit directly between vertebrate hosts and rely on tick bites for infection. More broadly, this work identifies a previously unrecognized host cell surface feature that limits viral entry and highlights the importance of monitoring viral evolution that could alter transmission potential.

## Introduction

Mammalian cells display a dense barrier at the cell membrane characterized by a complex network of glycans. Multiple viruses from different families use these glycans as attachment factors, including various types of glycosaminoglycans, N- and/or O-linked glycans on glycoproteins, and glycolipids [1–4]. Influenza viruses represent a paradigmatic example in which interaction with glycans is generally essential for viral entry. Specifically, classical influenza viruses attach to sialic acids (SAs), which are monosaccharide residues derived from neuraminic acid. Moreover, the linkage type between the SA and the terminal galactose of the glycoconjugate (α2,3 or α2,6) determines the avian or human tropism of influenza A viruses [4–7]. SAs can be present at the terminal end of N- or O-glycans, are abundantly displayed on the surface of many vertebrate cells, and are involved in the entry of other viruses, such as coronaviruses, reoviruses, and picornaviruses [1,5,8]. For instance, influenza C and D viruses, as well as embecoviruses such as OC43 coronavirus, bind 9-O-acetylated SAs [1,9–14].

The *Orthomyxoviridae* family comprises five genera in addition to influenza viruses [15–18], including three genera of fish viruses and two genera that infect terrestrial vertebrates, namely thogotoviruses and quaranjaviruses [19–23]. Thogotoviruses have gained prominence due to their ability to infect humans, with examples including Thogoto virus (THOV), Dhori virus (DHOV), Oz virus (OZV), and Bourbon virus (BRBV). For instance, in 2014 BRBV was associated with a human fatality in Kansas, and additional human cases have been reported since [19,20,24]. Anti-BRBV antibodies have also been detected in both wild and domestic animals [25,26]. OZV caused the death of a woman in Japan in 2023, and serological surveys suggest exposure in at least two additional individuals, as well as in wild mammals [22]. Human infections with THOV have been reported in Nigeria [27], and a DHOV infection occurred following accidental laboratory exposure [23]. Moreover, the detection of antibodies against THOV and DHOV in livestock and humans in several countries such as Spain and Portugal [28,29] highlights the significant circulation of these viruses and underscores the need to assess their zoonotic potential.

Thogoto and quaranjaviruses are arthropod-borne viruses, with ticks serving as the main vectors, although Sinu virus (SINUV), a phylogenetically divergent thogotovirus, was isolated from mosquitoes [30]. The vertebrate hosts of thogotoviruses include various mammalian species, such as rodents, dromedaries, and livestock including cattle [25,26,28,31]. Quaranjaviruses have been shown to possess a broad host range [18,21,32,33], and have also been occasionally detected in humans [21]. Notably, Wellfleet Bay virus (WBV) has caused recurrent mass mortality events among anseriforms in North America [32].

Like all orthomyxoviruses, thogoto and quaranjaviruses are enveloped viruses with a segmented, single-stranded, negative-sense RNA genome [19]. However, unlike influenza A and B viruses, they do not encode a neuraminidase. Instead, they express a single surface glycoprotein (GP) that is phylogenetically related to the GP64 protein of baculoviruses [34,35], linking them to other arthropod viruses. Viral entry occurs through endocytosis [20,34,36], but no entry receptors have been identified so far, and the role of SAs in viral entry remains unaddressed. A microglycan array using the BRBV GP revealed no hits, suggesting that glycans are not used for attachment or entry [37].

In this work, we aimed to better understand the influence of SAs on non-influenza orthomyxovirus entry. Using vesicular stomatitis virus (VSV) pseudotypes harboring different thogotovirus or quaranjavirus GPs, we show that SA removal or lectin binding to SAs significantly enhances viral entry for some representatives of these genera, and we confirmed this observation using authentic THOV and DHOV. Therefore, contrary to influenza viruses, SAs function as a barrier to GP-mediated viral entry. Finally, we performed experimental evolution of the SINUV and WBV GPs by passaging VSV recombinants in a human cell line that highly expresses SAs. We identified key mutations in the WBV GP that confer adaptation to an SA-enriched human cell culture at the entry stage, demonstrating the adaptive potential of these viruses to overcome the SA barrier to viral entry.

## Methods

### NCI60 panel data

The NCI-60 panel, which consists of 60 characterized cancer lines from various tissue origins, was obtained from the National Cancer Institute (dtp.cancer.gov/discovery_development/nci-60). We included 47 cell lines from this panel in our analyses. Additional information is provided in **Supplementary Data 1**. Processed RNA-seq expression data are available for each of these cell lines (discover.nci.nih.gov/cellminer/loadDownload.do; RNA-seq - composite expression) and are expressed as log_2_(FPKM) + 1), where FPKM are fragments per kilobase per million.

### Cells

SKMEL-28 and IGROV-1 cells from the NCI-60 panel were cultured in RPMI (Gibco) supplemented with 10% FBS (Gibco), 10 units/mL penicillin, 10 μg/mL streptomycin (Gibco), 250 ng/mL amphotericin B (Gibco) and 5 μg/mL prophylactic plasmocin (InvivoGen) at 37°C and 5% CO_2_. Vero and BHK-21 cells were cultured in DMEM, and HAEC primary cells in hAEC Culture Medium (Epithelix), in all cases at 37 °C with 5% CO_2_. For passaging, cells were resuspended with trypsin (Gibco), appropriately diluted, and transferred to new flasks. PCR assays regularly confirmed the absence of mycoplasma contamination.

### SA quantitation

Approximately 2.5 × 10^5^ living cells were spun down and washed with FACS buffer (PBS supplemented with 2% BSA). Biotinylated lectins MAL II (*Maackia Amurensis Lectin II*, L-1260-2, Vector labs) and SNA (*Sambucus Nigra Lectin*, B-1305-2, Vector labs) that specifically bind to α2,3- and α2,6-SAs, respectively, were added at 4°C for 30 min at a concentration of 20 µg/mL in FACS buffer. Then, lectin-treated cells were washed twice with FACS buffer and incubated with Streptavidin-Dylight488 (“strep-dye”, Vector labs, 1:1000 in FACS buffer) for 20 min in the dark at 4°C. Cells were washed twice and resuspended in 500 µL FACS buffer before being processed by flow cytometry to quantify mean fluorescence intensity. Cells treated with FACS buffer without lectins were used as a negative control and then incubated with strep-dye. Two technical replicates were carried out for each experiment.

### Viruses

We used THOV/SiAr/126/72 and DHOV/India/1313/6 isolates, derived from isolates 113.3 and 77.1, respectively [31]. These isolates were amplified in BHK-21 cells. A GFP-expressing Influenza A virus strain A/Puerto-Rico/8/34 (PR8) was kindly provided by Adolfo García-Sastre (Mount Sinai School of Medicine, New York).

### VSV pseudotype production

VSV pseudotypes harboring thogotovirus (SINUV, OZV) and quaranjavirus (WBV) GPs were produced in previous work [38]. Briefly, HEK293T cells were seeded in T75 flasks previously coated with poly-D-lysine (Gibco) and transfected the following day with 30 μg of pcDNA3.1-C-HisTag expression plasmids encoding codon-optimized GP sequences. At 24 h post-transfection, cells were transduced with GFP-encoding VSV particles lacking the cognate VSV G gene. Transduced cells were incubated for 24 h and pseudotypes were harvested by collecting the supernatant, cleared by centrifugation at 2000 g for 10 min, passed through a 0.45 μm filter, aliquoted, and stored at -80°C. Residual carryover VSV-G protein was neutralized with a monoclonal antibody obtained in house from a mouse hybridoma cell line. A VSV pseudotype bearing the VSV-G glycoprotein and a bald pseudotype carrying no added glycoprotein were produced to be used as controls.

### Infection with VSV pseudotypes

Cells were seeded in 96-well plates and, on the following day, culture medium was removed and cells were inoculated with VSV pseudotypes for 1 h at 37°C, after which RPMI 10% FBS was added. Viral entry was monitored after 18-24 h in an Incucyte SX5 Live-Cell Analysis System (Sartorius) by acquiring phase contrast and GFP confluence values. As a blank control of background signal due to remaining VSV-G carry-over or cell autofluorescence, cells were inoculated with the bald pseudotype. Two technical replicates were carried out for each assay. The percentage of infected cells was calculated as the ratio between GFP and phase contrast confluence, subtracting the corresponding blank control value.

### Lentiviral pseudotype production

Lentivirus pseudotypes were produced in previous work [38]. Briefly, HEK293T cells were seeded and transfected with pCMVΔR8.2 packaging plasmid, pTRIP-GFP plasmid and the same GP expression plasmids as above, incubated at 37°C and, 48 h post-transfection, supernatants were harvested, cleared by centrifugation at 2000 g for 10 min, aliquoted and stored at -80°C.

### Infection with lentiviral pseudotypes

Cells to be infected were seeded in 96-well plates the day before. At the time of infection, medium was removed from the cell culture and 50 µL of lentiviral pseudotype inoculum was added to each well. After 1 h at 37°C, 50 µL of RPMI 10% FBS was added to each well. Plates were incubated for two days at 37°C, after which they were imaged using the Incucyte SX5 Live-Cell Analysis System (Sartorius). The percentage of infected cells was calculated as the ratio between the GFP area and cell confluence.

### Growth curves with authentic THOV and DHOV

SK-MEL-28 cells were inoculated at a multiplicity of infection (moi) of 0.001 plaque forming units (pfu) per cell. After adsorption during 1.5h at 37°C, cultures were washed with PBS and DMEM supplemented with 1% FBS and antibiotics was added. Supernatants were collected at 0, 8, 12, 24, 48 and 72 hpi, and viral titers were determined as plaque forming units (pfu) per mL by plaque assays on Vero cells [39]. Viral growth curves were performed three times. Area Under the Curve (AUC) values were calculated using log_10_-transformed titers.

### Authentic THOV and DHOV plaque assays

SK-MEL-28 cells were seeded in 12-well plates and, the following day, culture medium was removed, cells were washed once with PBS and 200 µL of viral inoculum (several 10-fold dilutions, from 10^-2^ to 10^-6^) were added per well. After incubating infected cells during 1.5h at 37°C, the inoculum was removed and 1 mL of fresh culture medium containing 0.6% agar were added per well. After 72 h of incubation at 37°C, cells were fixed with 3.7% formaldehyde in PBS and stained with 0.1% crystal violet to visualize plaques.

### Neuraminidase treatment

Cells were seeded into a 96-well plate and treated with exogenous neuraminidase from *Clostridium perfringens* (exoNA) (Merck, N2876; 35mU/100 µL of culture medium), which removes SAs from glycan chains by cleaving their α2,3-, α2,6- or α2,8-linkages. Infections were carried out after overnight incubation also in the presence of exoNA. Non-exoNA-treated cells were used as a mock condition. Each virus and condition were assayed three times. For statistical analysis, the values representing the percentage of infected cells were log_10_-transformed by calculating log_10_(x+1), where x is the corresponding percentage of infected cells. The x + 1 term was added to prevent zeros from appearing in the logarithm calculation.

### Neuraminidase treatment of viral particles

Media containing viral particles were treated with exoNA for 2 h and centrifuged for 2 h at 30,000 g and 4°C to wash the exoNA. As a control, each virus was also subjected to the same procedure but without exoNA. The percentage of infected cells values were log_10_-transformed for statistical analysis.

### Lectin assays

Cells were pre-treated with MAL II and or SNA lectins diluted in Opti-MEM medium at the desired concentration (4, 20, 60 or 100 µg/mL) for 1 h at 37 °C before infection. For mock controls, cells were pre-treated with 50 µL per well of Opti-MEM without lectins. Inoculation of viruses was carried out as indicated above, and viral entry was monitored by fluorescence signal acquisition at 18-24 hpi at 37°C in an Incucyte SX5 Live-Cell Analysis System (Sartorius). Each virus and lectin condition were tested three times. The percentage of infected cells was log_10_-transformed by calculating log_10_(x+1), where x is the percentage of infected cells.

### Generation of GP-encoding VSV recombinants

Recombinant viruses were used for experimentally evolving the GPs. A cDNA VSV infectious clone encoding GFP and lacking the VSV-G gene was used to clone the GP sequence. To this end, expression plasmids encoding the wild-type GP sequence of SINUV and WBV (without codon optimization) were used for PCR using Phusion Hot Start II High-Fidelity PCR Master Mix (ThermoFisher). A pair of primers was used to amplify each GP (**Supplementary Data 2**). The PCR product was purified using the DNA Clean & Concentrator-5 kit (Zymo Research) and cloned with HiFi (NEBuilder) into the cDNA VSV backbone, using a pair of primers to open the backbone (**Supplementary Data 2**). The recovery of infectious particles from the cDNA infectious clone was carried out as detailed in previous work [40]. Briefly, BHK-G43 cells expressing the VSV-G glycoprotein through mifepristone induction were transfected with the cDNA clone along with helper plasmids encoding VSV N, P and L proteins and the T7 RNA polymerase using lipofectamine 3000 (Invitrogen). After 3 h at 37°C, the medium was replaced with DMEM 10% FBS supplemented with 10 nM mifepristone to induce VSV-G expression and cells were incubated at 33°C for 36 h, followed by 48 h at 37°C. Supernatants from GFP-positive cell cultures were harvested, clarified by centrifugation at 2000 g for 10 min, and used to inoculate non-VSV-G-expressing BHK-21 cells to amplify the virus. The viral inoculum was removed, and BHK-21 cells were washed five times with PBS and incubated in DMEM containing 2% FBS and 25% anti-VSV-G neutralizing monoclonal antibody. Supernatants were harvested at 24-48 hpi, clarified by centrifugation, aliquoted, and stored at -80°C.

### Experimental evolution of SINUV and WBV GP

SK-MEL-28 cells in 6-well plates were inoculated with GP-encoding recombinant VSV at an moi of 0.1 pfu/cell. Cells were incubated with the viral inoculum at 37°C, shaking every 15 min. After 1 h, cells were supplemented with DMEM containing 10% FBS and incubated at 37°C for 24 h to allow viral growth, after which supernatants were harvested, cleared by centrifugation at 2000 g for 10 min, aliquoted, and stored at -80°C. We performed 10 serial passages. Between each passage, supernatants were titrated on BHK-21 cells. Three independent evolutionary replicates were performed for each GP. Following the 10 serial passages, for all evolution replicates and for the parental GPs, viral growth curves were performed by infecting cells in the same manner as in the evolution passages, with or without exoNA treatment. GFP signal and cell confluence were acquired using an Incucyte SX5 Live-Cell Analysis System (Sartorius) at different time points of the kinetics.

### GP sequencing

Viral RNA was extracted from supernatants using the Quick-RNA Viral Kit (Zymo Research) following the manufacturer’s recommendations. SuperScript IV (Invitrogen) was used to reverse-transcribe the extracted viral RNA with a primer targeting a sequence in the VSV genome located upstream of the GP gene. PCR amplification of the GP gene was performed using Phusion Plus DNA Polymerase (Thermo Scientific) with specific primers targeting the flanking VSV genome sequences (**Supplementary Data 2**). PCR products were purified using the DNA Clean & Concentrator kit (Zymo Research) according to the manufacturer’s instructions and sequenced by linear amplicon sequencing (Eurofins Scientific) using the Oxford Nanopore technology.

### Viral GP alignments

GP amino acid sequences were retrieved from the National Center for Biotechnology Information (NCBI) RefSeq (www.ncbi.nlm.nih.gov/refseq) database. Sequences were aligned using Clustal Omega v1.2.4 (www.clustal.org/omega) and the alignment was visualized using the R package *ggmsa*. The following RefSeq accession numbers were included: YP_010840496.1 (SINUV), YP_009553284.1 (OZV), YP_010840292.1 (BRBV), YP_010840363.1 (UPOV), YP_010840326.1 (JOSV), YP_010840302.1 (TT-THOV), YP_009352874.1 (DHOV), YP_145808.1 (THOV), YP_010840455.1 (ARGV), YP_009508042.1 (QRFV), YP_009996582.1 (JAV), YP_009987463.1 (LCV), YP_010840351.1 (TLKV), and YP_009110688.1 (WBV).

### GP structural model

The SINUV and WBV GPs were folded with AlphaFold3 using AlphaFold Server (https://alphafoldserver.com). The trimer structure was obtained by specifying three copies of the protein in AlphaFold Server. To improve structural prediction, the first amino acids of the N-terminal region and the last amino acids of the C-terminal region were removed from the input protein sequence loaded into AlphaFold Server, as they corresponded to a disorganized region and the transmembrane region, respectively. For SINUV, this corresponded to the first 20 and last 50 residues, while for WBV this corresponded to the first 20 and last 48 residues.

### GP site-directed mutagenesis

GP expression plasmids were used for site-directed mutagenesis (Quickchange, Agilent) using PfuUltra High-Fidelity DNA Polymerase (Agilent) and a pair of overlapping primers containing the desired mutation (**Supplementary Data 2**), following the manufacturer’s instructions. Mutations were verified by Sanger sequencing of individual *E. coli* clones. The mutagenized expression plasmids were prepared and purified for pseudotype production following the protocol detailed above.

## Results

### Viral entry mediated by thogoto and quaranjavirus GPs negatively correlates with SA expression

In a previous study, we used the NCI-60 panel to characterize the cellular tropism of VSV pseudotypes carrying the envelope proteins of a panel of RNA viruses [38], including the GPs of WBV, SINUV, and OZV. Here, we repeated these assays with the GPs of the latter, adding as controls a GFP-expressing influenza A virus (PR8) and a VSV pseudotyped with the native glycoprotein. We then performed correlation analyses between infection rates and RNA-seq data for genes encoding sialyltransferases across the 47 NCI-60 cell lines for which such data were available. As expected, PR8 infectivity tended to correlate positively with sialyltransferase expression (**Figure 1A**). In contrast, entry mediated by the SINUV, OZV, and WBV GPs tended to correlate negatively with the expression of the sialyltransferases ST3GAL4, ST3GAL5, ST3GAL6, ST6GAL1, ST6GALNAC2, ST6GALNAC3, and ST8SIA1 (Pearson correlations, *P* < 0.01 in all cases).

**Figure 1.**
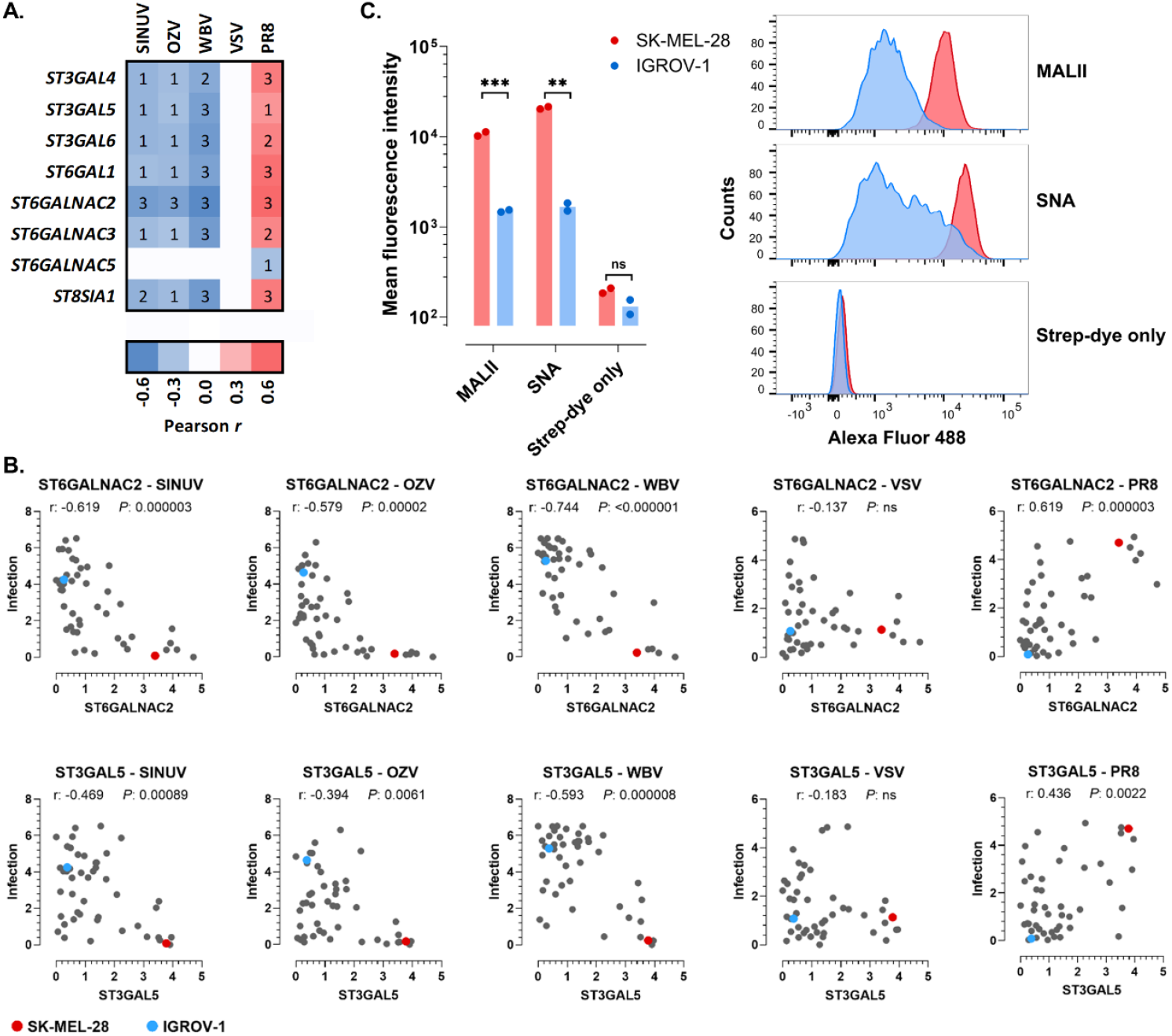
Negative association between SA content and GP-mediated viral entry. **A**. Correlations between RNA-seq data for sialyltransferases and infectivity of VSV pseudotypes harbouring WBV, SINUV or OZV GPs, as well as PR8 influenza. Gene expression data were quantified as fragments per kilobase per million (FPKM) and infection rate as the percentage of infected cells. Both metrics were scaled between 0 and 100, transformed as log_2_(x+1), and their Pearson correlation was calculated. Genes for which at least one virus showed a significant (*P* < 0.01) correlation are shown. The heatmap depicts the average Pearson correlation from two independent infection assays, as shown in the scale bar. The numbers inside the heatmap represent the levels of significance (1: *P* < 0.01, 2: *P* < 0.001, 3: *P* < 0.0001). VSV pseudotyped with its own glycoprotein was included as a control. **B**. Examples of scatter plots between infectivity and RNA-seq data for the ST6GALNAC2 and ST3GAL5 genes. Pearson correlation coefficients (r) and their statistical significance (*P*) are shown (ns: not significant). The data points for the SK-MEL-28 cell line are marked in red, and those for IGROV-1 cell line are marked in blue. **C**. SK-MEL-28 and IGROV-1 cells were incubated with biotin-labelled MAL II and SNA lectins to quantify by flow cytometry α2,3- and α2,6-linked SAs, respectively. Lectin-treated cells were incubated with Streptavidin-Dylight488TM (strep-dye) and the signal emitted by the dye was used to quantify SAs. Left: fluorescence data from two independent experiments. Cells treated without lectin and incubated with the strep-dye (strep-dye only) were used as negative control. A t-test between SK-MEL-28 and IGROV-1 values was carried out using log-transformed data (**: *P* < 0.01; ***: *P* < 0.001; ns: *P* > 0.05). Right: fluorescence level histogram from one representative experiment is shown. Y-axis shows the relative cell count (%) of α2,3- or α2,6-linked SAs-positive cells, and X-axis shows the intensity of fluorescence.

Among the cell lines in the NCI-60 panel, we aimed to identify those falling at the two extremes of the observed correlations and to directly quantify their SA content. To this end, we selected SK-MEL-28 cells (derived from a human melanoma) and IGROV-1 cells (derived from a human cystadenocarcinoma). SK-MEL-28 cells were among the least susceptible to thogoto/quaranjavirus pseudotypes yet were among the most highly infected by PR8, and they tended to show high sialyltransferase RNA-seq expression levels. In contrast, IGROV-1 cells showed the opposite pattern, that is, high susceptibility to thogoto/quaranjavirus pseudotypes, low susceptibility to PR8, and low sialyltransferase expression levels (**Figure 1B**). We then quantified the expression of α2,3- and α2,6-linked SAs in these two cell lines by flow cytometry. Staining with *Maackia amurensis* lectin II (MAL II) and *Sambucus nigra* lectin (SNA) indicated that SK-MEL-28 cells displayed approximately ten times more α2,3-/α2,6-linked SAs than IGROV-1 cells (t-test: *P* = 0.0008 for α2,3-linked SAs, *P* = 0.0017 for α2,6-linked SAs), consistent with the RNA-seq data (**Figure 1C**).

### Enzymatic SA depletion from cells enhances thogoto/quaranjavirus GP-mediated viral entry

We treated SK-MEL-28 cells with exogenous neuraminidase (exoNA) to reduce their SA content and then assessed changes in thogoto/quaranjavirus GP-mediated viral entry. We found that exoNA treatment strongly increased entry mediated by the SINUV GP (143-fold; t-test: *P* < 0.001) and the WBV GP (177-fold; *P* = 0.001; **Figure 2A**). In contrast, the same treatment strongly reduced PR8 infectivity (100-fold; *P* < 0.001). Significant but modest increases in infectivity were observed in control assays using VSV pseudotyped with its own glycoprotein (2.5-fold; *P* < 0.001). The increase observed for the OZV GP was similarly modest (5-fold; *P* < 0.001).

**Figure 2.**
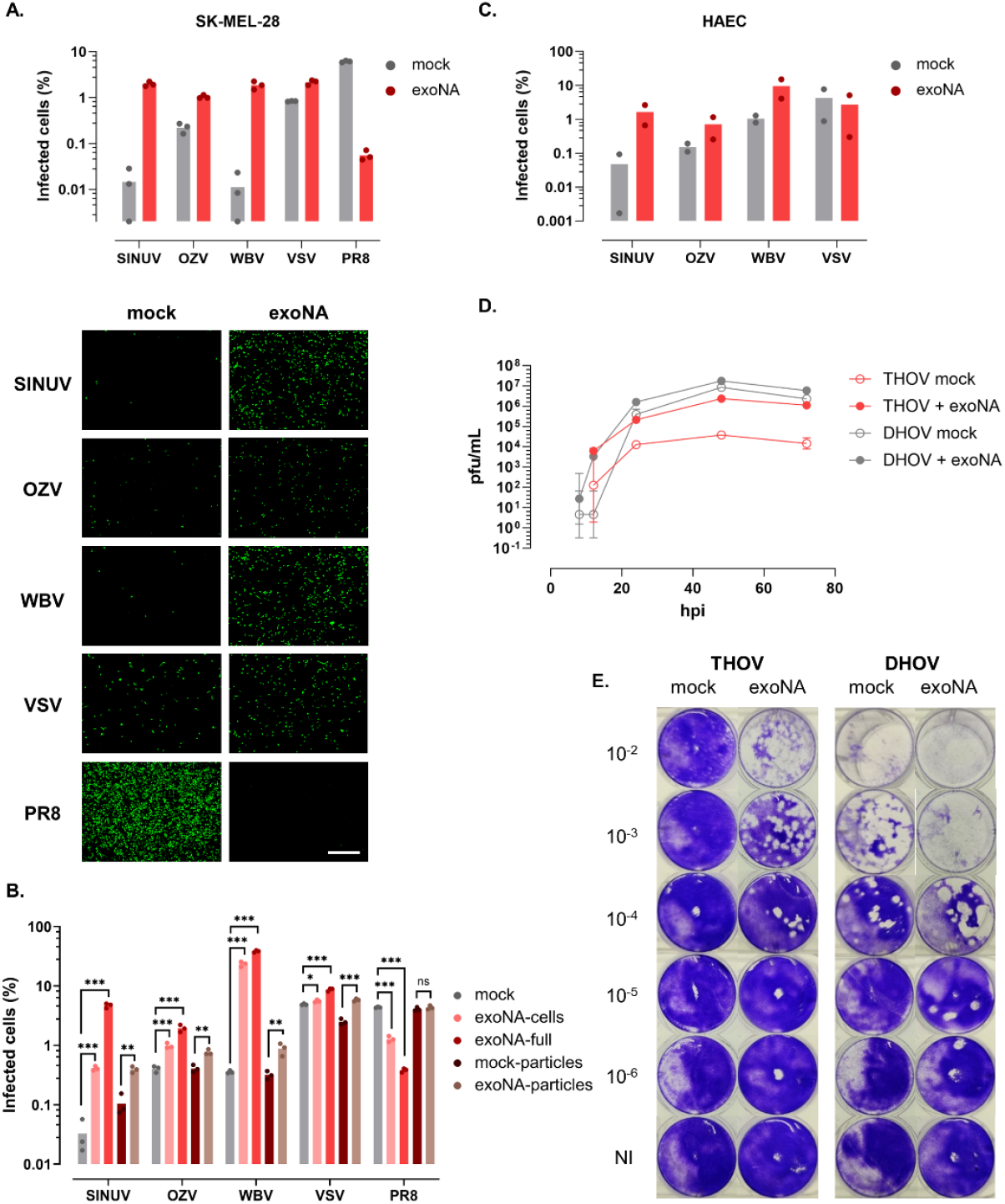
Effect of SA depletion on GP-mediated viral entry. **A**. Effect of exoNA treatment of SK-MEL-28 cells on the infectivity of VSV pseudotypes harbouring WBV, SINUV or OZV GPs, as well as PR8 influenza. VSV pseudotyped with its own glycoprotein was included as a control. Viral entry was measured as the percentage of GFP-positive cells at 18-24 hpi. Data points correspond to three independent experiments. Values at the bottom of the graph correspond to 0. Differences between mock and exoNA-treated cells were significant for the four viruses (t-tests: *P* < 0.001). Below: representative images (scale bar, 1000 µm). **B**. SK-MEL-28 cells were infected with the same viruses as above under five different conditions: no exoNA treatment (mock), exoNA pre-treatment of cells only (exoNA-cells), exoNA treatment in which cells were pre-treated and the exoNA was also present in the infection medium (exoNA-full), exoNA treatment of viral particles only (exoNA-particles), and control assay in which particles were processed in the same way but with no exoNA treatment (mock-particles). Data points from three independent experiments are shown. Each treatment condition was compared with its respective mock by performing a t-test using log-transformed data. *: *P* < 0.05; **: *P* < 0.01; ***: *P* < 0.001; ns: *P* > 0.05. **C**. Same assays as in panel A using HAEC primary cells. Data points from two independent experiments are shown. No statistical tests were performed since n = 2. **D**. Viral growth kinetics of authentic THOV and DHOV in SK-MEL-28 cells in the presence or absence of exoNA treatment (moi = 0.001). Supernatants were collected at different time points and titrated in Vero cells. Means and standard deviations of viral titers (pfu/mL) from three independent experiments are shown for each time point. **E**. Plaque forming efficiency of THOV and DHOV in SK-MEL-28 cells in the presence or absence of exoNA. Several 10-fold dilutions of each virus were used as viral inoculum (from 10^-2^ to 10^-6^) and culture medium containing 0.6% agar was added. 72 hpi cells were fixed with formaldehyde and stained with crystal violet. A non-infected condition is also shown (NI).

While these results suggest that SAs on the cell surface can function as a barrier to thogoto/quaranjavirus entry, a possible confounder of the above assays was that exoNA was still present during infection, meaning that SAs from viral particles could also be digested. To address this, we repeated these assays by pre-treating SK-MEL-28 cells with exoNA and performing the infection assays after exoNA removal. The effects on thogoto/quaranjavirus GP-mediated viral entry were smaller but overall similar to those observed with the full treatment (SINUV GP: 13-fold, OZV GP: 3-fold, WBV GP: 66-fold; t-tests: *P* < 0.001 in all cases; **Figure 2B**). The slightly reduced effect could be explained by the recovery of SAs during infection. Indeed, PR8 infectivity was also mildly affected by this treatment (4-fold; *P* < 0.001). In parallel, we treated viral particles with exoNA, washed them, and inoculated cells with these pre-treated particles. We found smaller enhancements of infectivity under these conditions (less than 4-fold in all cases), confirming that the observed effects of exoNA were mainly mediated by depletion of cellular SAs.

We then tested whether our results were specific to the SK-MEL-28 cell line. For this purpose, we used primary human airway epithelial cells (HAEC). Despite some cytotoxicity following exoNA treatment, manifested as a reduction in cellular confluence, we still observed enhanced viral entry (**Figure 2C**). Specifically, exoNA treatment increased the infectivity of VSV pseudotypes harboring the SINUV, OZV, and WBV GPs by 35-fold, 5-fold, and 9-fold, respectively. Again, no such effect was observed in control assays using VSV pseudotyped with its own glycoprotein.

We also sought to determine whether the observed effects were specific to the VSV pseudotyping system. To address this, we produced lentiviral pseudotypes harboring the WBV GP. This experiment confirmed that exoNA treatment of SK-MEL-28 cells strongly enhanced WBV-mediated viral entry. Specifically, the percentage of infected cells increased from 0.03% under control conditions to 1.02% following SA removal, representing a 30-fold increase (t-test: *P* = 0.003).

Based on these results, we aimed to confirm our observations using available authentic viruses, for which we used THOV and DHOV, each representing one of the two main phylogroups within the *Thogotovirus* genus [31]. Viral growth curves (**Figure 2D**) performed on SK-MEL-28 demonstrated a strongly positive effect of exoNA treatment on THOV titers, as determined by plaque assays in Vero cells. For example, at 48 hpi, THOV reached 2.4 × 10^6^ pfu/mL in exoNA-treated cells, versus 4.0 × 10^4^ pfu/mL in untreated cells (60-fold effect). For DHOV, we observed an initial acceleration of growth, such that at 12 hpi, viral growth was detectable only in exoNA-treated cells. However, at later time points, differences in titer became smaller (ca. 3-fold). Overall, the area under the curve (AUC) was significantly higher in exoNA-treated cells for both THOV (t-test: *P* < 0.001) and DHOV (*P* = 0.006).

We also tested for changes in plaque forming efficiency by performing plaque assays of a viral stock in SK-MEL-28 cells. A dramatic effect was observed for THOV, which was able to form obvious plaques only in the exoNA-treated cells (**Figure 2E**). In contrast, DHOV was able to form large plaques either in the presence or absence of exoNA, albeit the cytopathic effect appeared to be slightly higher in exoNA-treated cells. Overall, the experiments with authentic viruses confirmed the effects of SA depletion observed with the pseudotypes of closely related thogotoviruses, and suggested differences between members of the *Thogotovirus* genus.

### Lectins enhance thogoto/quaranjavirus GP-mediated viral entry

As an alternative approach to digesting SAs with exoNA, we set out to incubate cells with lectins, which should bind SAs and make them less accessible to viral particles. We again used MALII and SNA, which specifically bind to α2,3- and α2,6-linked SA, respectively. PR8 infectivity was modestly reduced after incubating SK-MEL-28 cells with these lectins (**Figure 3**). For instance, treatment with 100 µg/mL of SNA lectin reduced the number of PR8-positive cells by approximately two-fold (t-test: *P* > 0.05), whereas MALII has little or no effect at all. The combined treatment with both lectins had similar effects as SNA alone. We interpret that the observed effects should result from the competition between lectins and viral particles for access to SAs.

**Figure 3.**
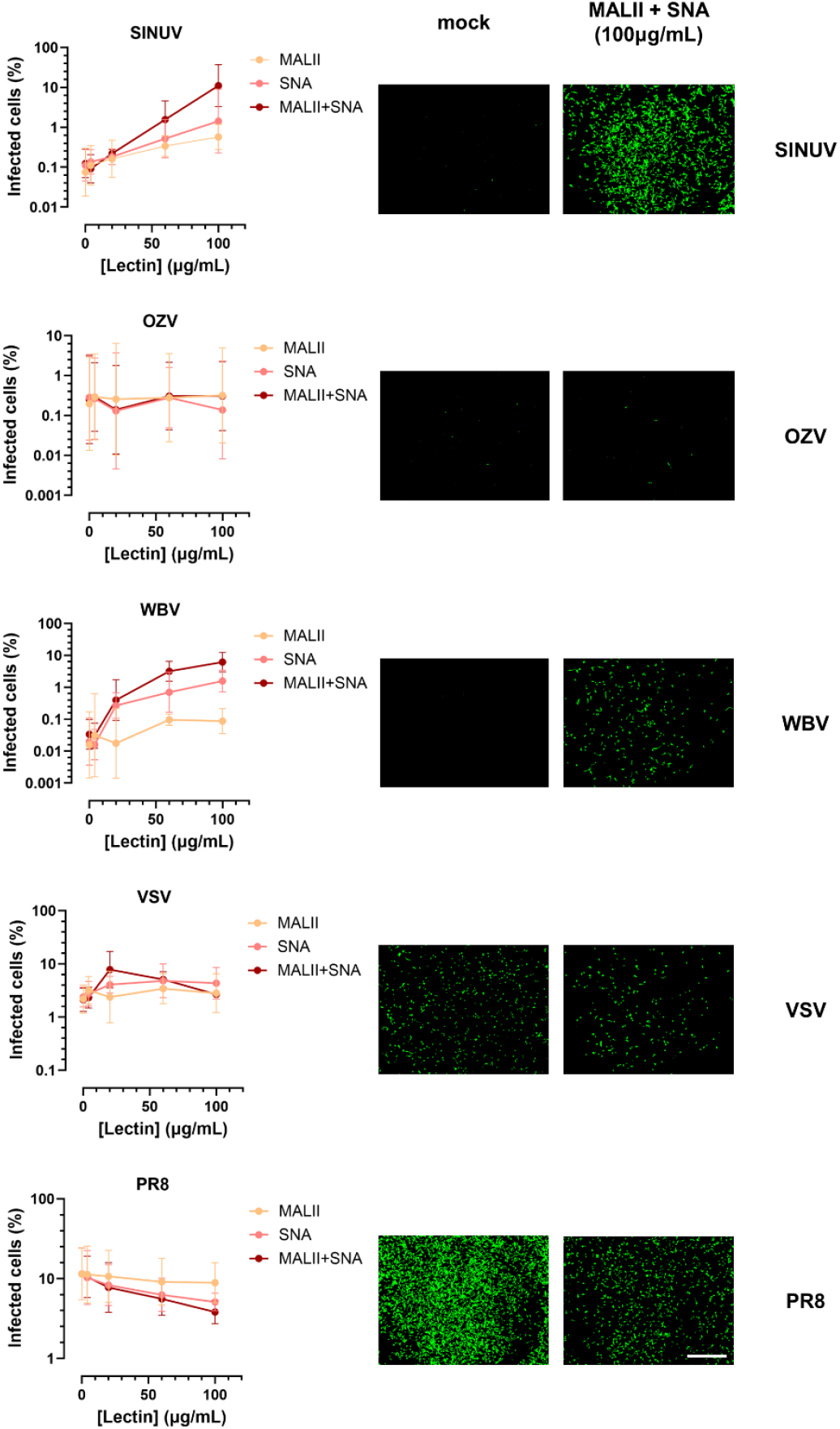
Effect of SA-specific lectins on GP-mediated viral entry. Effect of MALII (α2,3-SA specific) and SNA (α2,6-SA specific) lectins on the infectivity of VSV pseudotypes harbouring WBV, SINUV or OZV GPs, as well as PR8 influenza. VSV pseudotyped with its own glycoprotein was included as a control. Viral entry was measured as the percentage of GFP-positive cells at 18-24 hpi. Three independent assays were carried out for each lectin dose. Right: representative images showing the effects of the combined MALII and SNA treatment at a dose of 100 μg/mL each (scale bar, 1000 µm). Statistical tests comparing the infectivity data at 0 versus 100 μg/mL are provided in the text.

In contrast to the results obtained for PR8, a strongly dose-dependent effect on infectivity was observed for VSV pseudotypes harbouring the GPs of SINUV and WBV. Notably, the combined treatment with MALII and SNA at 100 µg/mL increased infectivity by >100-fold in both cases (t-tests: *P* = 0.018 and *P* = 0.005, respectively). When only one lectin was added, the positive effects on infectivity were less strong than with the combined treatment, but still stronger for SNA (100 µg/mL: 20-fold for SINUV, t-test: *P* = 0.111; 50-fold for WBV, *P* = 0.031) than for MALII (100 µg/mL: 6-fold for SINUV and 3-fold for WBV; *P* > 0.05). In contrast, lectin treatments had no effect on VSV pseudotyped with its own glycoprotein or with OZV GP. Overall, these observations recapitulate the results obtained using exoNA.

### Experimental evolution of SINUV and WBV GPs

To assess the ability of thogoto and quaranjavirus GPs to overcome the barrier to viral entry imposed by SAs, we engineered replication-competent recombinant VSVs encoding the GPs of WBV or SINUV, and serially passaged them in SK-MEL-28 cells. We selected these two GPs because they showed the most evident response to exoNA and lectin treatments in the above experiments with VSV pseudotypes. We avoided evolving authentic viruses due to gain-of-function concerns. For each GP, three independent evolution lines were maintained for 10 passages (**Figure 4A**). Then, the founder and evolved viruses were evaluated by performing growth curves in the presence or absence of exoNA. Confirming the above results, viral growth/replication of the recombinant viruses were higher in exoNA-treated cells, but this enhancement became less marked in the evolved viruses (**Figure 4B**). Hence, in passage-10 viruses, the course of infection in exoNA-treated cells more closely resembled the mock-treated condition than it did in founder viruses. This suggests that the GPs may have adapted to bypass the restrictions on viral entry imposed by SAs. However, this shift was more obvious for VSV-WBV than for VSV-SINUV.

**Figure 4.**
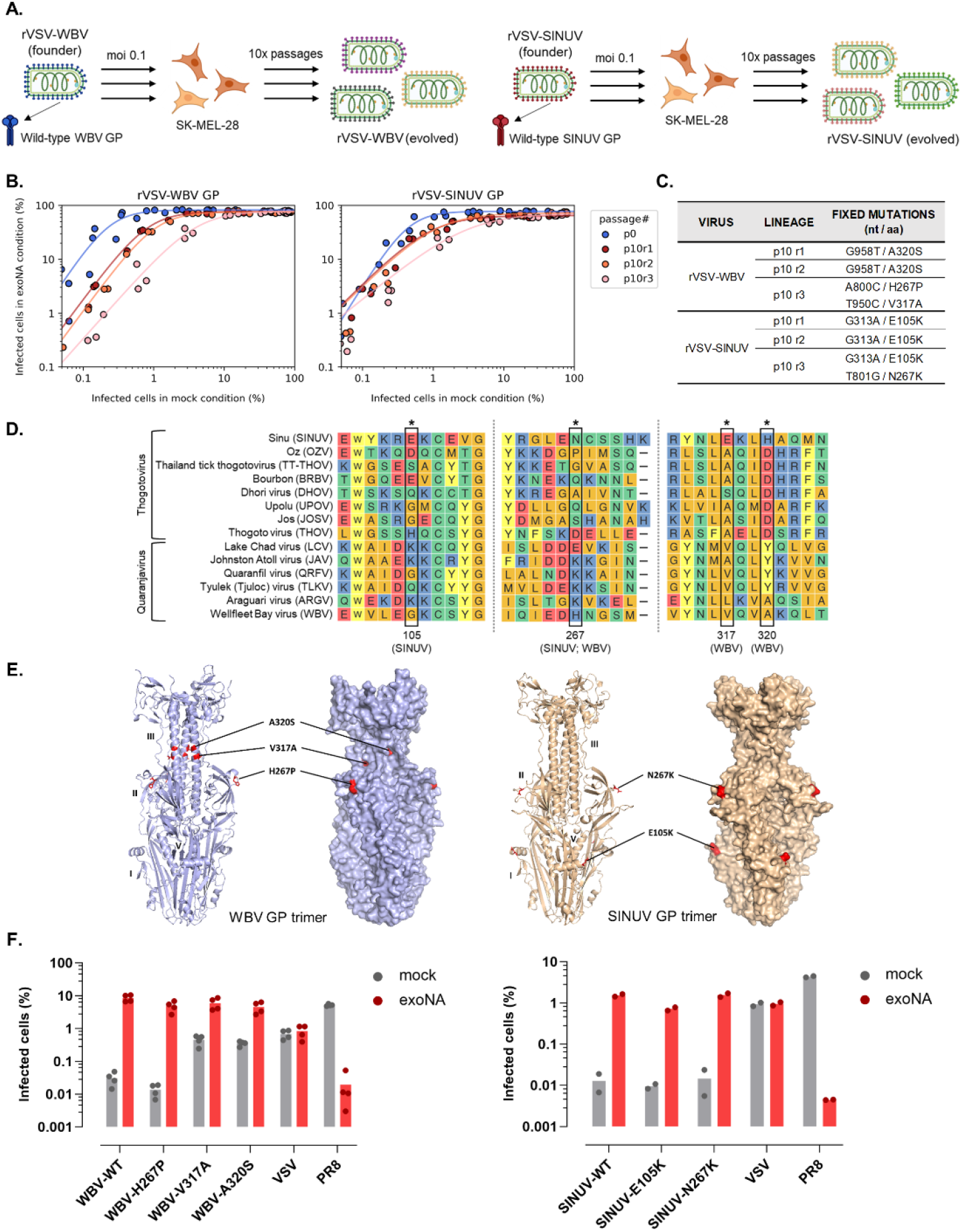
Adaptation of SINUV and WBV GPs to SK-MEL-28 cells. **A**. Scheme of the evolution experiment. **B**. Representation of viral growth curves in exoNA-treated cells relative to the mock condition. The percentage of infected cells is shown. Each data point corresponds to a growth curve time point. Data are represented as a function of the mock condition instead of as a function of time to correct for differences in growth kinetics resulting from adaptations unrelated to SAs (e.g. faster VSV replication). Founder viruses are shown in blue, and the three evolved viruses in shades of red. Lines indicate a logistic growth model fit to the data by least-squares regression (goodness of fit: r^2^ > 0.95 in all cases). **C**. Amino acid exchanges in the evolved and founder GPs. **D**. Clustal omega alignment of thogoto and quaranjavirus GP sequences. Only the positions that showed substitutions in our experimental evolution and their flanking regions are shown. **E**. Structural models of the SINUV and WBV glycoprotein trimer predicted by AlphaFold3. The exchanged positions are indicated in red. Some of the glycoprotein domains are indicated in the structures. **F**. Effects of exoNA treatment on the infectivity of VSV pseudotypes harboring WBV or SINUV GPs with the parental sequence or each assayed mutant. Individual data points from four independent experiments for WBV GP and from two independent experiments for SINUV GP are shown. Doses of pseudotypes were adjusted to obtain similar infectivity values among GP variants in the exoNA condition. Statistical tests comparing the effect of the exoNA treatment between GP variants are provided in the text.

To elucidate the genetic basis of these phenotypic changes, we sequenced the evolved viral GPs from passage 10. Three fixed amino acid substitutions were found in the WBV GP (H267P, V317A, A320S), and two were found in the SINU GP (E105K, N267K; **Figure 4C-D**). Notably, the A320S substitution appeared in two of the three evolved WBV GPs, while the E105K substitution appeared in all three evolved SINUV GPs. Moreover, position 267 independently evolved in WBV and SINUV GPs, albeit the identity of the parental and evolved residues differed. These cases of parallel evolution strongly suggest that the observed substitutions were adaptive, although their effects may not be related to SAs. In structural models of the GP trimers from SINU and WBV obtained with AlphaFold3, the SINUV N267K and the WBV H267P substitutions were found to be located in structurally equivalent areas at the GP surface (**Figure 4E**). The E105K substitution was also located at the surface of the GP trimer. The V317A and A320S mutations, on the other hand, mapped to a less-exposed GP region.

To more directly evaluate the effects of each mutation in relation to SAs, we introduced them individually in GP-expression plasmids, produced VSV pseudotypes, and assayed their infectivity in SK-MEL-28 in the presence or absence of exoNA. In these assays, we adjusted titers such that infectivity values were similar across GP variants under the more permissive exoNA-treated condition. We found that the fold change in infectivity in the absence of exoNA compared to exoNA-treated cells was similarly strong for the parental and H267P WBV GPs (276-fold and 355-fold, respectively; t-test comparing log fold-increases: *P* > 0.5; **Figure 4F**). However, the effects of the exoNA treatment were less marked for the V317A and A320S WBV GP mutants (13-fold for both mutants, t-tests comparing with the log-fold increase of the WT: *P* < 0.001). For SINUV, the effect of exoNA treatment was 120-fold for the parental GP, versus 76-fold for E105K and 106-fold for N267K, revealing no significant differences among these GP variants. Therefore, the response of the SINUV GP to exoNA treatment did not evolve so markedly as for the WBV GP, consistent with the growth curves performed with recombinant VSVs.

## Discussion

Here, we aimed to explore the relationship between SAs and viral entry in human-infective orthomyxoviruses outside the influenza genus. The results were unexpected, as we found that SAs play opposite roles in SA-binding influenza A and quaranja and thogoto orthomyxoviruses. However, not all thogoto and quaranjavirus GPs responded in the same way to the presence of SAs, with SINUV and WBV showing the greatest sensitivity. SINUV is a divergent relative of thogotoviruses, while WBV is a quaranjavirus. We also note that THOV and DHOV, which belong to different thogotovirus subclusters, might differ in their SA sensitivities.

We confirmed this negative impact of SAs on thogoto and quaranjavirus entry using pseudotypes for both viruses and authentic thogotoviruses. While the use of authentic viruses is important to validate the relevance of our results, pseudotyped viruses present several advantages, as they allow us to focus specifically on entry and to work with a broader range of viruses, including high-containment or non-culturable viruses. Moreover, pseudotypes allow the use of wild-type sequences, ensuring that the interpretation of results is not confounded by tissue-culture adaptation. In this context, it is worth noting that, since their original isolation from ticks [41,42], THOV and DHOV have been amplified in mammalian cells and thus undergone replication in the presence of SAs.

While SAs have been extensively characterized as attachment factors promoting viral entry, fewer studies have examined their role as entry barriers. One possible mechanism by which SAs may hamper viral entry is by acting as decoy receptors [8]. This has been demonstrated for influenza A virus and mucosal SAs, where sialylated compounds in respiratory mucus can trap virions and prevent infection [43–45]. In this context, the virus-encoded neuraminidase plays a key role by cleaving mucosal SAs, allowing viral particles to be released and potentially engage SAs on cell surface proteins. A similar process occurs in embecoviruses such as human coronavirus OC43, which encode a hemagglutinin-esterase functionally analogous to the influenza neuraminidase [46]. Whether the decoy mechanism applies to thogoto and quaranjaviruses remains to be demonstrated. Previous work failed to identify glycans that bind to the BRBV GP [37], arguing against this hypothesis. Additionally, unlike influenzaviruses and embecoviruses, thogoto- and quaranjaviruses possess no known neuraminidase or hemagglutinin-esterase activities. On the other hand, it has been observed that THOV GP can induce hemagglutination, although the specific glycan responsible for this process remains unknown [47].

An alternative mechanism explaining the observed barrier effect could involve shielding of the entry receptor, or specific residues within it, by SAs. This is analogous to findings with SARS-CoV-2, where SAs associated with ACE2 modestly interfere with spike binding [48]. However, our results with lectins do not support this hypothesis, as lectins, which would be expected to further sterically hinder virus–receptor interactions, in fact strongly promoted viral entry in some cases. Testing this hypothesis would be greatly facilitated by the identification of thogoto and quaranjavirus entry receptors, which remain currently unknown.

Human infections with some thogoto and quaranjaviruses have been reported, underscoring their zoonotic potential [19–23]. However, no direct viral transmission has been found among vertebrate hosts, as infection occurs exclusively through tick bites [49–51]. Since mucosa are particularly rich in SAs, our results suggest that SAs may represent a significant barrier to thogoto and quaranjavirus transmission via mucosal routes, such as airborne, sexual, or orofecal pathways. According to this hypothesis, direct injection of the virus via tick bites would efficiently bypass the mucosal SA barrier, leading to successful infection of vertebrates. Transmission from the bloodmeal into cells of the tick gut may not be subject to these restrictions, as the quantity and composition of SAs in ticks differs from those of vertebrates [52–54].

Finally, our work reveals the ability of WBV GP to adapt to the presence of SAs, illustrating the evolutionary potential of these viruses to overcome the obstacle posed by SAs. Overcoming the SA barrier could enhance inter-mammalian including human-to-human transmissibility. Our experiments explored how point mutations could reduce the sensitivity of GPs to SAs. Other detected mutations do not appear to be related to SAs, such as the mutation at position 267. This position is mutated in both SINUV and WBV GPs. In SINUV, it is found within a glycosylation motif (Asn-X-Ser/Thr; X ≠ Proline), whereas in WBV it is adjacent to another residue also located inside a glycosylation motif. Mutations at this site may therefore lead to the loss of glycosylation sites, potentially contributing to GP optimization under the experimental conditions used here.

The neuraminidase of influenza viruses not only facilitates the release of progeny virions form the cell, but also enhances the mobility of influenza A virus through the SA rich mucus layer of the respiratory tract [43–45,55]. It is conceivable, that yet another potential evolutionary outcome might be acquisition of a neuraminidase function from influenza viruses. This would require coinfection of the same host and stable incorporation of the neuraminidase-encoding segment into thogoto or quaranjavirus particles. The likelihood of a reassortment event between quaranjaviruses and influenzaviruses is difficult to estimate, but this possibility may be of concern for WBV, which infects aquatic birds, increasing the chances of contact with influenza viruses.

## Acknowledgments

We thank I. Andreu-Moreno and R. Martínez-Recio for technical assistance. This work was financially supported by a European Research Council Advanced Grant (101019724—EVADER) and a grant from the Spanish Ministerio de Ciencia e Innovación (PID2020-118602RB-I00— ZooVir) to R.S. J.M.-G. is funded by a PhD fellowship from the Spanish Ministerio de Ciencia, Innovación y Universidades (FPU fellowship FPU21/03807). J.D. is the recipient of an European Molecular Biology Organization postdoctoral fellowship (ALTF 140-2021) and a Marie Skłodowska-Curie Actions Postdoctoral Fellowship (101104880). The funders had no role in study design, data collection and analysis, decision to publish or preparation of the manuscript.

